# Salient sounds distort time perception and production

**DOI:** 10.1101/2022.07.04.498704

**Authors:** Ashley Symons, Fred Dick, Adam Tierney

## Abstract

The auditory world is often cacophonous, with some sounds capturing attention and distracting us from our goals. Despite the universality of this experience, many questions remain about how and why sound captures attention, how rapidly behavior is disrupted, and how long this interference lasts. Here we use a novel measure of behavioral disruption to test two predictions made by models of auditory salience. First, models predict that goal-directed behavior is disrupted at points in time that feature a high degree of spectrotemporal change. We find that behavioral disruption is precisely time-locked to the onset of distracting sound events: participants tapping to a metronome speed up immediately after the onset of distractors. Moreover, this response is greater for more salient sounds (larger amplitude) and sound changes (greater pitch shift). Second, models predict that different auditory features are combined into an overarching salience map. We find that the time course of behavioral disruption is highly similar after acoustically disparate sound events, suggesting overlapping mechanisms of attentional capture: both sound onsets and pitch shifts of continuous background sounds speed responses at 750 ms, with these effects dying out by 1750 ms. These temporal distortions can be observed using only data from the first trial across participants. A potential mechanism underlying these results is that arousal increases after distracting sound events, leading to an expansion of time perception, and causing participants to misjudge when their next movement should begin.

**Significance Statement:** The noisy world constantly challenges our pursuit of goals. When driving, for example, a cacophony of mechanical, musical, and conversational sounds surrounds us and can wrench our focus away from the road. While the physiological signatures of auditory attentional capture are well researched, we know surprisingly little about how sound affects moment-to-moment behavior: How quickly do sounds affect our actions, how transient is the effect, and how is action affected by changes in sound properties? Here we use a synchronized tapping paradigm to show that loud sounds and large acoustic changes cause rapid distortions in time perception. However, these distortions were corrected within 2 seconds, showing that goal-directed behavior is transiently vulnerable yet ultimately resilient in the face of distraction.

## Introduction

Acoustic environments are complex, presenting a steady stream of interruptions that can interfere with goal-directed behavior. In a coffee shop, for example, you must focus on clearly communicating your order, while ignoring a dozen conversations, traffic noise, the hiss of the espresso machine, and miscellaneous electronic hums. Although distraction is a common experience, we know surprisingly little about how sound affects ongoing behavior: what factors cause a sound to disrupt behavior, how rapid is this disruption, and how long does the interference last?

In the visual system, researchers can track eye movements to measure attentional capture by parts of a visual scene (1). Using eye tracking data as ground truth, researchers have built computational models of visual salience in which several feature maps with center-surround inhibition are combined to form a “salience map” (2). Researchers studying auditory salience have created similar maps in time-frequency “space” (3-6). An alternate approach is to track feature-specific deviance from prediction relative to local and longer-term statistics (7-8).

These auditory salience models were validated by relating model predictions to subjective ratings (3-5, 7-11). Research has also investigated effects of sound presentation on performance on a difficult unrelated task, such as short-term memory (12-15) or perceptual detection (16). This research demonstrated that auditory scenes and objects featuring a greater degree of spectrotemporal modulation were rated as more salient and interfered with task performance to a greater degree. However, because these techniques cannot precisely measure the time course of behavioral disruption, it has not been possible to test several key predictions made by auditory salience models. First, they predict that ongoing goal-directed behavior will be disrupted at points in time that feature a high degree of spectrotemporal modulation. To test this prediction, behavioral disruption measurements must be precisely time-locked to sound events. Second, they predict that different auditory features are combined into an overarching saliency map, and that therefore changes in different dimensions (such as amplitude and pitch) disrupt behavior via the same mechanism. This prediction could be tested by examining the time course of behavioral disruption: if different auditory features capture attention via overlapping mechanisms, then the latency and duration of the disruption should be similar across dimensions.

Some prior evidence supporting these predictions comes from research on physiological components of the orienting response (17). The onset of loud sounds, for example, leads to time-locked changes in pupil dilation and the galvanic skin response, compared to the onset of soft sounds (18-20; but see 21). Sudden shifts in frequency have also been linked to changes in pupil size, with greater dilation for larger shifts (22-24). Stimuli with greater high-frequency amplitude modulation (roughness) have been linked to microsaccadic inhibition (11). Degree of spectrotemporal modulation in task-irrelevant distractor sounds has also been linked to the magnitude of EEG effects, including increased P3a amplitude (25-26) and decreased phase-locking and gamma responses to task-relevant sounds (27). Unlike the behavioral measures of attentional capture reviewed above, the orienting response is precisely time-locked to stimulus features; however, it not yet been clearly linked to behavioral disruption. Based on these physiological studies, we hypothesize that sudden acoustic changes are rapidly followed by a disruption of ongoing behavior, but that individuals can rapidly recover from this interference.

We tested the prediction that task-irrelevant sound onsets and pitch changes lead to a rapid, transient disruption in ongoing behavior by asking participants to tap to a click track, with differentially salient distractor sounds or sound changes occasionally presented between clicks. This task samples behavioral disruption at the tapping rate (2 Hz), enabling investigation of its time course. We predicted that distractor presentation would rapidly but briefly disrupt performance, pulling participants’ taps away from the click track. Further, this disruption should be greater for more salient distractors.

## Results

We conducted six experiments on short-term attentional capture. Due to the COVID-19 pandemic, three experiments were initially conducted online, with three subsequent matched in-lab replications. In each experiment, participants tapped to a click track at a rate of 2 Hz, attempting to align their movements with the clicks as precisely as possible. Occasionally, a 200-ms distractor was presented, with an onset exactly halfway between clicks (250 ms after the previous click). Attentional capture was measured as the change in the asynchrony between the participant’s tap and the nearest click for the four time points following the distractor. Given that the click rate was 2 Hz, this means that attentional capture was sampled at 250, 750, 1250, and 1750 ms after distractor presentation.

In Experiments 1 (online) and 2 (in-lab) we measured the effect of distractor roughness on attentional capture (Figure 1). Distractors were white noise carriers that were manipulated to be either more rough (100% amplitude modulated at 60 Hz) or less rough (20% amplitude modulated at 60 Hz). Collapsing across the more rough and less rough conditions, in both experiments we found an initial shift towards earlier tapping (online, at 250 ms, z = -5.10, p_corrected_ < 0.001; in-lab, at 750 ms, z = -3.11, p_corrected_ = 0.007), followed by a slowing down of tapping at 1250 ms (online, z = 4.53, p_corrected_ < 0.001; in-lab, z = 3.98, p_corrected_ < 0.001). There was, however, no difference between the more and less rough distractors at any time point (p_corrected_ > 0.05).

**Figure 1.**
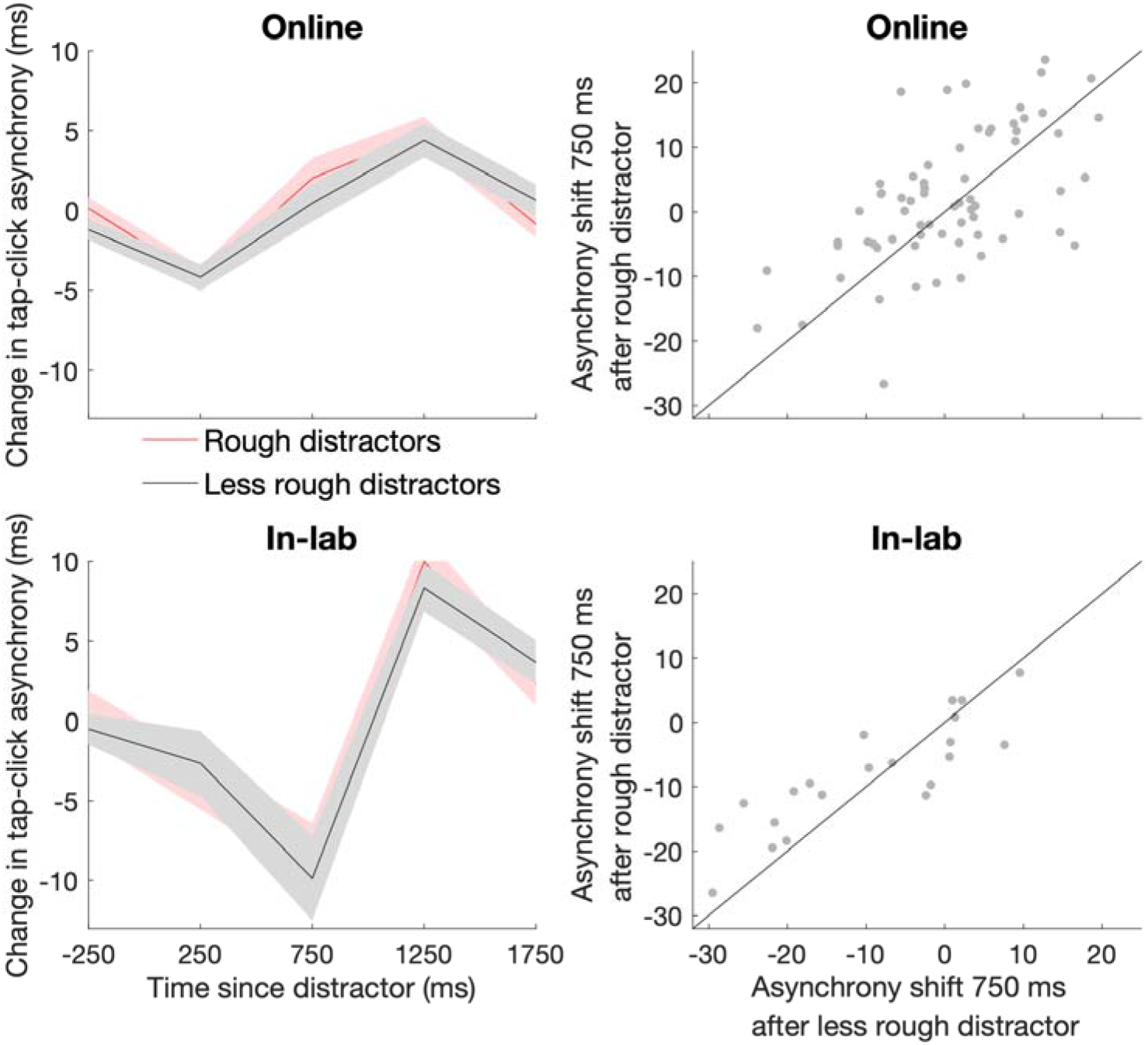
Effects of task-irrelevant sounds on auditory-motor timing are not modulated by distractor roughness. (Left) Mean change in tap-click asynchrony after distractor presentation in online study (top) and in-lab replication (bottom). Distractors were either rough (100% modulation at 60 Hz, red lines) or less rough (20% modulation at 60 Hz, black lines). The shaded region indicates the standard error of the mean. (Right) Scatterplot displaying relationship between the asynchrony shift after rough and less rough distractors in online study (top) and in-lab replication (bottom). The line displays the identity function y = x.

In Experiments 3 (online) and 4 (in-lab) we measured the effect of distractor volume on attentional capture (Figure 2). Distractors were a drill sound that was presented at a loud or soft volume (rms dB difference = 19.17). Collapsing across the loud and soft conditions, in both experiments we found an initial shift towards earlier tapping (Wilcoxon signed rank test; online, at 750 ms, z = -2.78, p_corrected_ = 0.022; in-lab, at 250 ms, z = -3.66, p_corrected_ = 0.001), followed by a slowing down of tapping at 1250 ms (online, z = 3.06, p_corrected_ = 0.009; in-lab, z = 3.92, p_corrected_ < 0.001). In both experiments the loud distractor led to earlier tapping at 750 ms compared to the soft distractor (online, z = -3.24, p_corrected_ = 0.005; in-lab, z = -3.25, p_corrected_ = 0.005). This difference, though reliable, was small in magnitude, with a median 4.6 ms difference between conditions in both Experiment 3 and Experiment 4. Only in Experiment 4 was tapping later at 1250 ms for the loud distractor (z = 2.58, p_corrected_ = 0.040).

**Figure 2.**
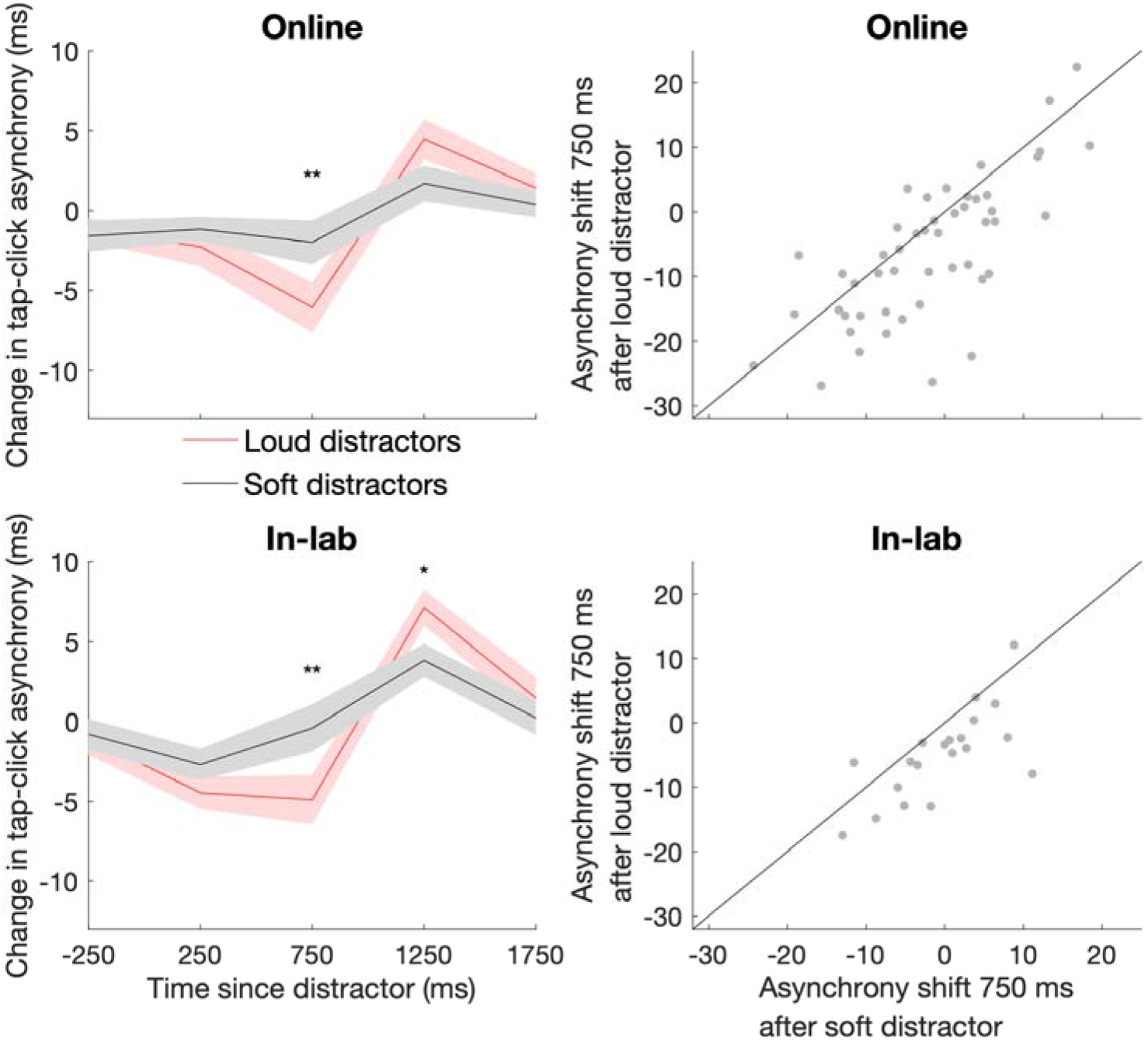
Effects of task-irrelevant sounds on auditory-motor timing are modulated by distractor volume. (Left) Mean change in tap-click asynchrony after distractor presentation in online study (top) and in-lab replication (bottom). Distractors were either loud (red lines) or soft (black lines; rms dB difference = 19.17). ** indicates time points at which p_corrected_ < 0.01, * indicates time points at which p_corrected_ < 0.05. The shaded region indicates the standard error of the mean. (Right) Scatterplot displaying relationship between the asynchrony shift after soft and loud distractors in online study (top) and in-lab replication (bottom). The line displays the identity function y = x.

In Experiments 5 (online) and 6 (in-lab) we measured the effect of the magnitude of pitch change on attentional capture (Figure 3). Alongside the click track we presented a stream of tones at a rate of 20 Hz. Distracting sound events consisted of a brief change in the pitch of the tone stream which lasted for 200 ms; this change was either large (6 semitones) or small (1 semitone). (Importantly, although 1 semitone is a small change relative to the large change condition, it is well above most individuals’ detection threshold (28).) Collapsing across the large and small pitch change conditions, we found no overall effect of distractor presentation on tapping asynchrony (all p_corrected_ > 0.05). However, in both experiments the large pitch shift led to earlier tapping at 750 ms compared to the small pitch shift (online, median difference 2.1 ms, z = -3.29, p_corrected_ = 0.004; in-lab, median difference 6.2 ms, z = -2.90, p_corrected_ = 0.015).

**Figure 3.**
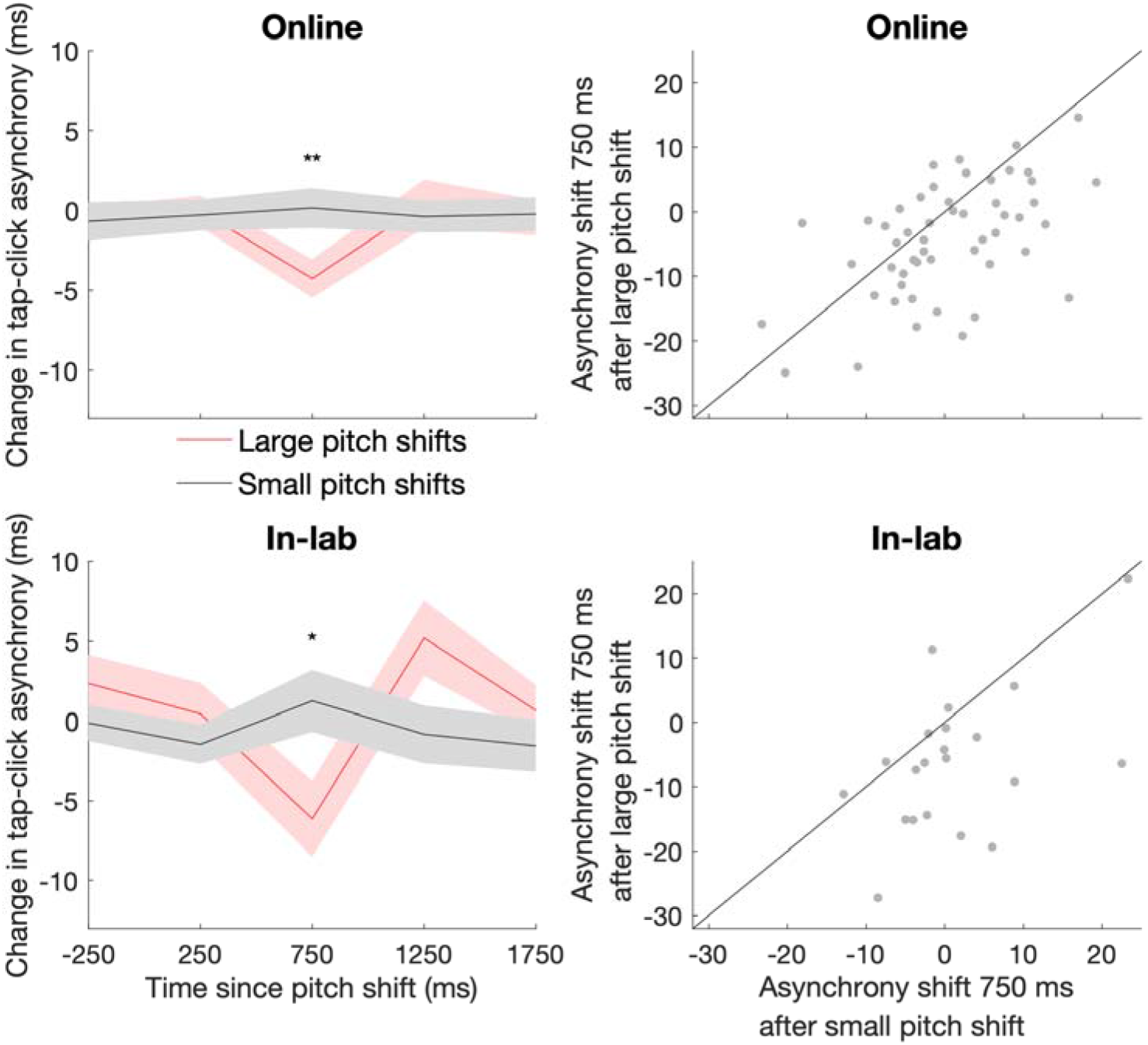
Effects of task-irrelevant pitch changes on auditory-motor timing are modulated by change magnitude. (Left) Mean change in tap-click asynchrony after change in pitch of a constant background sound in online study (top) and in-lab replication (bottom). Pitch changes were either large (6 st, red lines) or small (1 st, black lines). ** indicates time points at which p_corrected_ < 0.01, * indicates time points at which p_corrected_ < 0.05. The shaded region indicates the standard error of the mean. (Right) Scatterplot displaying relationship between the asynchrony shift after large and small pitch changes in online study (top) and in-lab replication (bottom). The line displays the identity function y = x.

Overall, findings were broadly replicable across the online and in-lab experiments: distractor presentation was linked to an initial shift towards earlier tapping followed by a decrease in tempo, with the magnitude of the initial shift modulated by volume and pitch shift magnitude but not roughness. However, although clear tapping shifts were visible for most individual participants (Figure S1), there was considerable variability across participants in the timing and extent of tapping shifts. To determine whether there existed reliable individual differences in attentional capture, for each experiment we used Spearman’s correlations to compare the size of the tapping shift at each time point across the two conditions. Across all six experiments, the resulting correlations were strongest at 750 ms, and so we focus on this time point here. Cross-condition correlations at 750 ms were, for Experiment 1, rho = 0.56, p < 0.001; for Experiment 2, rho = 0.88, p < 0.001; for Experiment 3, rho = 0.70, p < 0.001; for Experiment 4, rho = 0.73, p < 0.001; for Experiment 5, rho = 0.51, p < 0.001; and for Experiment 6, rho = 0.39, p = 0.081.

Across multiple experiments, the effect of the distractor on tapping synchronization could be observed within a single trial (Figure 4). Collapsing across conditions, we tested whether there was a reliable change in asynchrony across participants following the first presentation of the distractor in each experiment. In Experiments 1 and 2, there was a shift towards earlier tapping on the first trial at 750 ms (Wilcoxon signed rank test; online, z = -4.660, p_corrected_ < 0.001; in-lab, z = -3.136, p_corrected_ = 0.003) followed by a slowing down of tapping (online, 1250 ms, z = 3.284, p_corrected_ = 0.001; in-lab, 1750 ms, z = 3.136, p_corrected_ = 0.003). In Experiments 3 and 4, there was a shift towards earlier tapping on the first trial at 750 ms in the online (z = -2.989, p_corrected_ = 0.011) but not in-lab experiment (z = -1.941, p_corrected_ > 0.05). In Experiments 5 and 6, there was a shift towards earlier tapping following the first pitch change at 750 ms in the in-lab experiment (z = -2.520, p_corrected_ = 0.047) but not in the online experiment (z = -0.046, p_corrected_ > 0.05).

**Figure 4.**
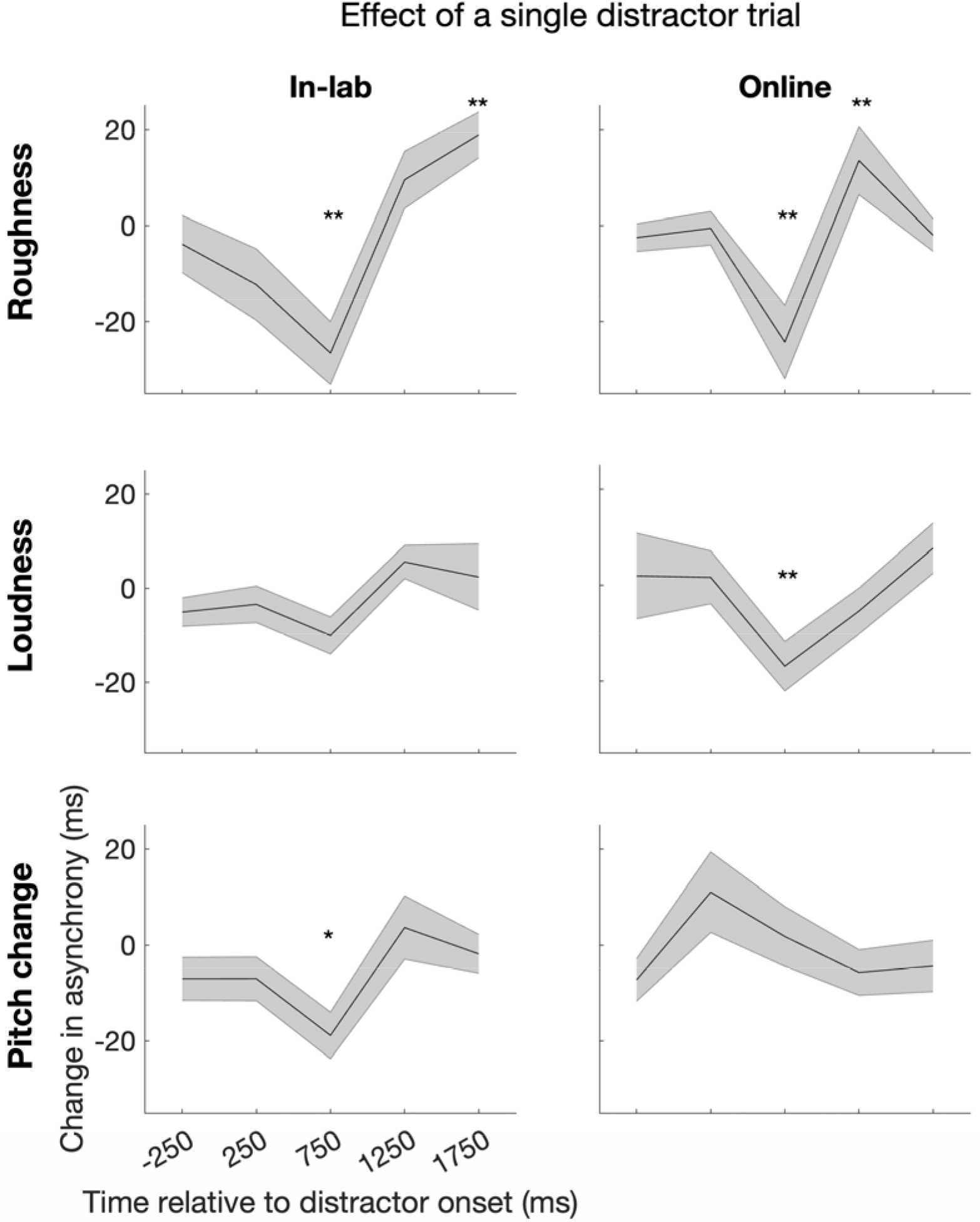
Effects of the first distractor on auditory-motor timing across roughness (top), loudness (middle), and pitch change (bottom) experiments. Each plot shows the mean change in tap-click asynchrony after distractor onset in the online experiments (left) and in-lab replication (right). ** indicates time points at which p_corrected_ < 0.01, * indicates time points at which p_corrected_ < 0.05. The shaded region indicates the standard error of the mean.

## Discussion

Our results confirm two key hypotheses made by computational models of auditory salience (3-8). First, essentially all models of auditory salience predict that acoustic edges will be salient. Here, we show that acoustic onsets and pitch changes cause rapid, transient disruption in ongoing synchronized tapping behavior, with greater effects for more salient sounds. Second, models of auditory salience suggest different features are combined into a single overarching salience map. We find that the task disruption across experiments follows a highly similar time course, with effects of sound volume and pitch shift size appearing at 750 ms and no longer evident by 1750 ms, suggesting that different types of sound events disrupt behavior via overlapping mechanisms. The time course of the performance impairment also roughly matches prior reports of physiological and electroencephalographic effects of salient sound presentation, suggesting a link between the orienting response and disruption of goal-directed behavior. For example, neural encoding of an attended tone stream decreases during the 2.3 seconds following the occurrence of a salient event in an unattended auditory scene (27), and pupil dilation responses modulated by sound volume are largely limited to the first two seconds following sound presentation (19).

Not only was synchronization impaired following distractor presentation, but it was biased: participants’ tapping shifted to be earlier in time. This suggests that they over-estimated the passage of time, mis-judging when the next click was about the arrive and therefore planning to move too early. What mechanism could explain this bias? One possibility is that salient sound presentation leads to an increase in arousal, speeding up the rate of internal pacemakers (29). Prior research has found that time perception expands after experimental manipulations designed to increase arousal, including modulation of body temperature (30), presentation of click trains (31-33), flickering of visual stimuli (34-35), and emotional intensity (36). In these studies time perception was measured using behavioral judgments which required participants to encode, remember, and compare temporal intervals to standards. Interestingly, the degree of temporal distortion we report here (1% of the interval being timed) is far lower than that reported in previous studies of temporal distortion due to manipulation of arousal. Presentation of click trains, for example, can distort time by as much as 10% (31). This discrepancy in effect magnitude between the current and previous findings suggests that arousal may distort time perception at both encoding and later stages such as retrieval, with distortion at later stages possibly being more severe than at earlier stages.

An alternate explanation for our findings is that synchronization to a metronome requires neural entrainment to the target rhythm (37), and that presentation of a distractor in between clicks briefly interferes with rhythmic temporal expectation (38). This explanation could account for the transient nature of the distractor effect, as the constant reinforcement of the metronome beat would enable participants to rapidly re-entrain after perturbation. This account, however, would have difficulty accounting for the fact that participants’ tapping consistently shifted to be earlier. Since distractors were presented exactly halfway between clicks, an entrainment perturbation account would have no reason to predict that tapping would shift in one direction versus the other.

Individual differences in the magnitude of timing shifts due to distractor presentation were highly reliable, with cross-condition correlations reaching as high as 0.88 in one experiment. This suggests that there are large, stable differences between participants in the extent of attentional capture due to presentation of task-irrelevant sounds. The source of these individual differences is an interesting target for future research. One possibility, for example, is that participants with greater inhibitory control may be better able to inhibit the capture of attention by task-irrelevant sounds.

Auditory roughness is a common feature of natural alarm signals (39), and previous reports have demonstrated a relationship between auditory roughness and both salience ratings and microsaccadic inhibition (11). Nevertheless, we did not find a significant effect of roughness on attentional capture. One possible explanation is that both the high roughness and low roughness stimuli presented in these experiments began with an aversive sharp, wide-band increase in amplitude. This sudden spectrotemporal change may have disrupted behavir robustly enough that the consequences of the subsequent amplitude modulation could not be detected due to ceiling effects.

The measure of behavioral disruption presented here has considerable methodological advantages over other similar measures. A single highly reliable measurement can be collected in 3-4 minutes. The task can be completed online, with similar results from online and in-lab experiments. A significant effect can be captured in a single trial, making possible analysis of changes in salience over time (for example, resulting from stimulus repetition). The task is simple and measures a natural behavior (40); as a result, it can be performed by virtually anyone over the age of 4 (41). The task does not rely on assessment of performance as correct or incorrect, and therefore is not susceptible to ceiling or floor effects. Finally, the task is a direct measure of behavioral disruption, and unlike salience ratings does not rely on participants’ interpretation of task instructions. This measure, therefore, could be an ideal tool for resolving conflicting predictions made by competing theories of auditory salience. Theoretical models of auditory salience, for example, differ on whether salience is primarily driven by local center-surround contrast (3-6) versus tracking of statistics on a longer time scale (7-8). This issue could be investigated by determining whether the temporal distortion response can be suppressed when task-irrelevant sounds are fully predictable.

## Materials and Methods

### Participants

All online participants were recruited through Prolific. The experiment was conducted via the online experiment platform Gorilla and all participants were asked to wear headphones. Automated procedures ensured that participants were using Google Chrome web browser on a computer. The Ethics Committee in the Department of Psychological Sciences at Birkbeck, University of London approved all experimental procedures. Informed consent was obtained from all participants. Participants were compensated for their participation in the form of payment at a standard rate. For each participant, trials in which the maximum tapping asynchrony was more than +/-250 ms were excluded. For online studies, participants with fewer than 36(/42) remaining trials were excluded from analysis.

101 participants (39 female, 61 male) between the ages of 18-40 (mean age = 24.19, sd = 5.79) took part in Experiment 1, with a final sample of 67 participants (29 female, 37 male, mean age = 24.61, sd = 6.01). 21 participants (10 female, 11 male) between the ages of 22-52 (mean age = 33, sd = 7.13) took part in Experiment 2. 62 participants (19 female, 43 male) between the ages of 18-52 (mean age = 26.03, sd = 6.82) took part in Experiment 3, with a final sample of 49 participants (16 female, 33 male, mean age = 26.69, sd = 7.21). 20 participants (3 female, 16 male, 1 other gender) between the ages of 20 – 42 (mean age = 31.65, SD = 7.67) took part in Experiment 4. 58 participants (28 female, 29 male, 1 other gender) between the ages of 18-52 (mean age = 26.90, sd = 6.13) took part in Experiment 5, with a final sample of 52 participants (26 female, 25 male, mean age = 26.92, sd = 6.28). The same participants who took part in Experiment 2 also took part in Experiment 6.

### Stimuli

In Experiments 3 and 5, stimuli consisted of 42 2-Hz isochronous click sequences that were each 8 seconds in duration, with distractor sounds presented in one of 7 positions (after the 6th through 12th clicks for Experiment 1 and after the 5th through 11th clicks for Experiment 3). In Experiments 1, 2, 4, and 6, stimuli consisted of 2 157.5-second 2-Hz isochronous click sequences, each of which contained 40 distractor sounds pseudo-randomly located such that there were at least 6 clicks between successive distractor sounds. There were 10 clicks at the start of the stimulus and 6 clicks at the end after which no distractor occurred. For Experiments 2, 4, and 6 stimuli were presented using PsychToolbox (version 3.0.17) run in Matlab (MathWorks, Inc) and the sound was delivered via Sennheiser HD 25-1 ii headphones.

In each experiment distractors were 200 ms in duration with an onset beginning 250 after the previous click. In Experiments 1 and 2, distractors were white noise carriers with a 10-ms cosine on/off ramp and amplitude modulated deeply (100%) or shallowly (20%) at 60 Hz. In Experiments 3 and 4, distractors were a pneumatic drill sound (Zhao et al., 2019) which was presented at a loud versus soft amplitude (rms dB difference = 19.17). In Experiments 5 and 6, alongside the click track a tone sequence was presented at a rate of 20 Hz. The 50-ms pure tones (10-ms on/off ramp) in the sequence had a fundamental frequency of 440 Hz and had a maximum amplitude of 0.6 relative to the clicks. Within each tone sequence there was a pitch deviation of either +1 semitone or +6 semitones that lasted for 200 ms (i.e. four 50-ms tones). The amplitude of the pitch deviation tones was reduced to balance the relative loudness of the tones using the stationaryLoudness function in Matlab (Stephen Hales Swift, 2020).

### Procedure

For Experiments 1, 3, and 5, participants were provided with on-screen task instructions asking them to tap along with the clicks by pressing the ‘j’ key on the keyboard while ignoring a distracting sound. Participants were provided with an example of the click sequence without the distractor so that they could practice tapping along to the beat. For Experiments 2, 4, and 6, participants were provided with verbal task instructions asking them to tap on a microphone along with the clicks. Experiments 3 and 5 lasted approximately 20 minutes, while Experiments 1, 2, 4, and 6 lasted approximately 7-8 minutes.

### Data Processing and Analysis

For Experiments 1, 3, and 5, sound timing information and participant response times were automatically recorded. For Experiments 2, 4, and 6, a stereo file was recorded with the stimulus in one channel and the participants’ response in the other channel. Custom Matlab scripts detected click and tap onsets by setting a threshold and relaxation time. Onsets were marked when the amplitude exceeded threshold and the amount of time elapsed since the last onset exceeded the relaxation time. Thresholds and relaxation times were adjusted on a participant-by-participant basis to ensure that each click and tap was marked and that no onsets were erroneously detected.

For each click, the difference between the participant’s tap time and the time of click onset was recorded. For Experiments 1, 3, and 5, the true asynchrony between tap times and click onsets could not be measured reliably, due to variations in sound onset latency resulting from differences in computer setups across participants. To create a measure of timing which was reliable and valid across both online and in-lab experiments, therefore, we measured change in tap-click asynchrony by calculating the difference between the asynchrony at each time point and the asynchrony at the previous time point. This procedure normalizes any cross-participant or cross-trial differences in latency. Change in asynchrony was assessed for the four time points following distractor presentation. To determine whether there was a significant overall change in asynchrony, Wilcoxon signed rank tests were conducted comparing the change in asynchrony to zero after collapsing across conditions. Wilcoxon signed rank tests were also used to examine whether the magnitude of the change in asynchrony was modulated by distractor parameters. P-values were Bonferroni corrected, given that comparisons were run across 4 time points. To examine performance on the first trial, data from the first trial was extracted and Wilcoxon signed rank tests used to determine whether there was a change in overall asynchrony relative to zero at each time point. One participant in Experiment 2 and one participant in Experiment 6 who tapped at antiphase following the first distractor was excluded from this analysis. For all experiments, only time points with significant effects are reported (for all other time points, p_corrected_ > 0.05). Statistics for all comparisons can be found in the Supplementary Materials. Processed data are available at https://osf.io/x8nhm/.

## Acknowledgements

The authors thank the participants. The authors are grateful to Clare Press for her helpful comments on an earlier version of the manuscript. This research is funded by the National Institutes of Health (NIH R01DC017734), the Leverhulme Trust (RPG-2019-107), and the Economic and Social Research Council (ES/V007955/1). None of the authors have potential conflicts of interest to disclose.

