## Supplementary Materials for "Salient sounds distort time perception and production"

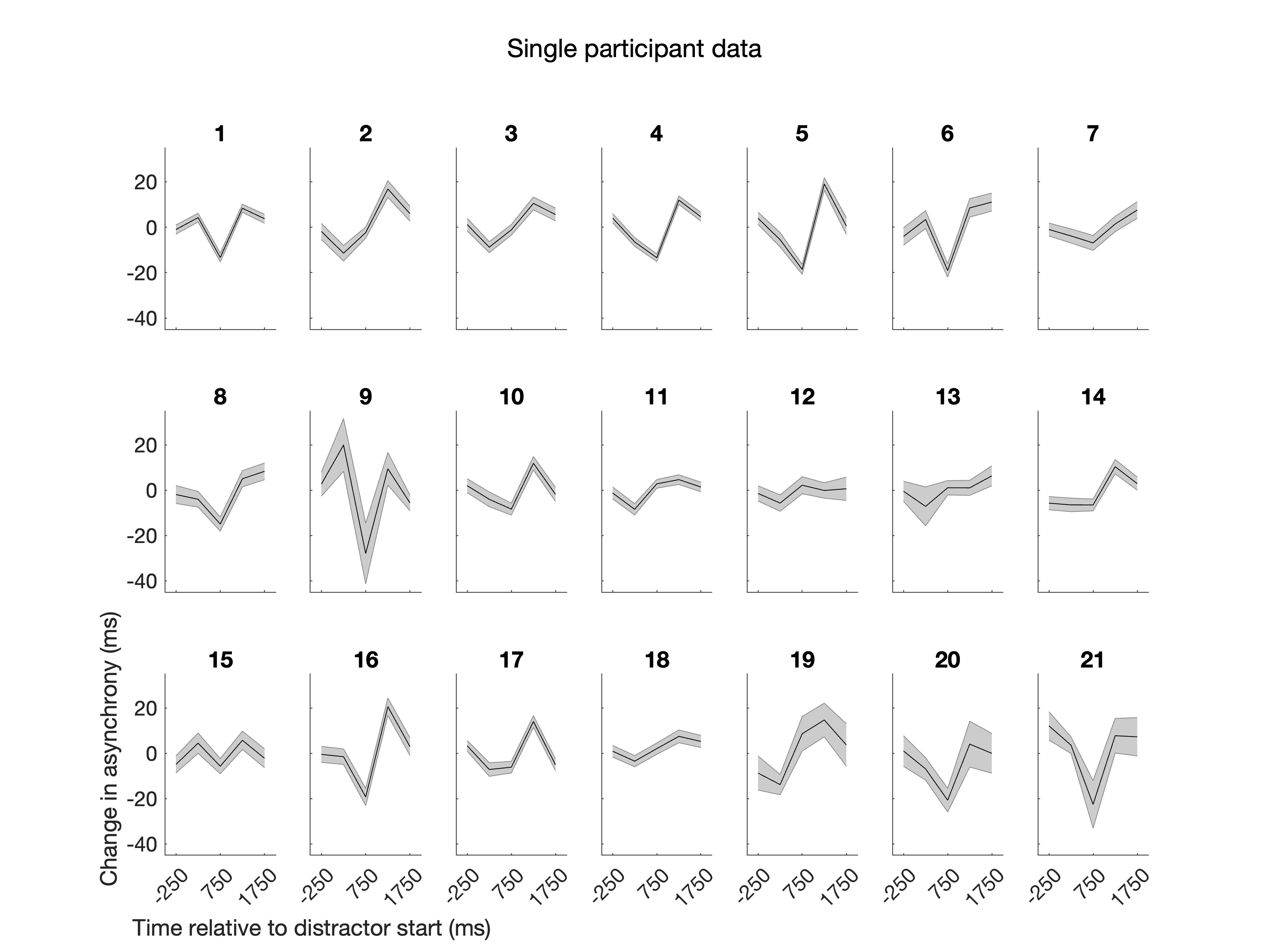

**Figure S1.** Effect of task-irrelevant sounds on audio-motor timing in single participants. Mean changes in tap-click asynchrony after distractor presentation are displayed for each of the 21 participants in Experiment 2. The shaded region indicates the standard error of the mean across trials.

**Table 1**. One-sample Wilcoxon tests comparing the change in tapping asynchrony in each condition and time point to zero for Experiments 1 and 2.

|  | Distractor Type | Time Point | B | W | z | p | p (corrected) |
| --- | --- | --- | --- | --- | --- | --- | --- |
| Experiment 1 (online) | High roughness | 250 | 67 | 415 | -4.523 | 0.000 | 0.000 |
|  |  | 750 | 67 | 1351 | 1.324 | 0.186 | 0.298 |
|  |  | 1250 | 67 | 1836 | 4.354 | 0.000 | 0.000 |
|  |  | 1750 | 67 | 994 | -0.906 | 0.367 | 0.489 |
|  | Low roughness | 250 | 67 | 360 | -4.866 | 0.000 | 0.000 |
|  |  | 750 | 67 | 1206 | 0.419 | 0.678 | 0.775 |
|  |  | 1250 | 67 | 1724 | 3.654 | 0.000 | 0.001 |
|  |  | 1750 | 67 | 1174 | 0.219 | 0.829 | 0.829 |
| Experiment 2 (in-lab) | High roughness | 250 | 21 | 48 | -2.346 | 0.018 | 0.023 |
|  |  | 750 | 21 | 20 | -3.319 | 0.000 | 0.001 |
|  |  | 1250 | 21 | 229 | 3.945 | 0.000 | 0.000 |
|  |  | 1750 | 21 | 158 | 1.477 | 0.147 | 0.147 |
|  | Low roughness | 250 | 21 | 62 | -1.860 | 0.065 | 0.074 |
|  |  | 750 | 21 | 35 | -2.798 | 0.004 | 0.007 |
|  |  | 1250 | 21 | 218 | 3.563 | 0.000 | 0.000 |
|  |  | 1750 | 21 | 183 | 2.346 | 0.018 | 0.023 |

**Table 2**. Wilcoxon signed rank tests comparing the change in tapping asynchrony between high and low roughness conditions at each time point for Experiments 1 and 2.

|  | Time Point | N | W | z | p | p (corrected) |
| --- | --- | --- | --- | --- | --- | --- |
| Experiment 1 (online) | 250 | 67 | 1175 | 0.2249 | 0.825 | 0.825 |
|  | 750 | 67 | 1401 | 1.637 | 0.102 | 0.408 |
|  | 1250 | 67 | 1197 | 0.3623 | 0.719 | 0.825 |
|  | 1750 | 67 | 1001 | -0.862 | 0.39 | 0.78 |
| Experiment 2 (in-lab) | 250 | 21 | 94 | -0.7473 | 0.473 | 0.785 |
|  | 750 | 21 | 150 | 1.199 | 0.243 | 0.785 |
|  | 1250 | 21 | 127 | 0.3997 | 0.708 | 0.785 |
|  | 1750 | 21 | 107 | -0.2954 | 0.785 | 0.785 |

**Table 3**. One-sample Wilcoxon tests comparing the change in tapping asynchrony in each condition and time point to zero for Experiments 3 and 4.

|  | Distractor Type | Time Point | N | W | z | p | p (corrected) |
| --- | --- | --- | --- | --- | --- | --- | --- |
| Experiment 3 (online) | Loud distractor | 250 | 49 | 411 | -2.004 | 0.045 | 0.12 |
|  |  | 750 | 49 | 266 | -3.447 | 0 | 0.002 |
|  |  | 1250 | 49 | 988 | 3.735 | 0 | 0.001 |
|  |  | 1750 | 49 | 726 | 1.415 | 0.158 | 0.181 |
|  | Soft distractor | 250 | 49 | 452 | -1.597 | 0.112 | 0.181 |
|  |  | 750 | 49 | 469 | -1.427 | 0.156 | 0.181 |
|  |  | 1250 | 49 | 765 | 1.517 | 0.131 | 0.181 |
|  |  | 1750 | 49 | 675 | 0.6217 | 0.541 | 0.541 |
| Experiment 4 (in-lab) | Loud distractor | 250 | 20 | 16 | -3.323 | 0 | 0.001 |
|  |  | 750 | 20 | 30 | -2.8 | 0.004 | 0.007 |
|  |  | 1250 | 20 | 202 | 3.621 | 0 | 0 |
|  |  | 1750 | 20 | 135 | 1.12 | 0.277 | 0.369 |
|  | Soft distractor | 250 | 20 | 44 | -2.277 | 0.021 | 0.034 |
|  |  | 750 | 20 | 99 | -0.224 | 0.841 | 0.869 |
|  |  | 1250 | 20 | 188 | 3.099 | 0.001 | 0.003 |
|  |  | 1750 | 20 | 110 | 0.1867 | 0.869 | 0.869 |

**Table 4**. Wilcoxon signed rank tests comparing the change in tapping asynchrony between loud and soft distractors at each time point for Experiments 3 and 4.

|  | Time Point | N | W | z | p | p (corrected) |
| --- | --- | --- | --- | --- | --- | --- |
| Experiment 3 (online) | 250 | 49 | 493 | -1.189 | 0.239 | 0.239 |
|  | 750 | 49 | 287 | -3.238 | 0.001 | 0.004 |
|  | 1250 | 49 | 860 | 2.462 | 0.013 | 0.026 |
|  | 1750 | 49 | 744 | 1.308 | 0.194 | 0.239 |
| Experiment 4 (in-lab) | 250 | 20 | 77 | -1.045 | 0.312 | 0.416 |
|  | 750 | 20 | 18 | -3.248 | 0 | 0.002 |
|  | 1250 | 20 | 174 | 2.576 | 0.008 | 0.017 |
|  | 1750 | 20 | 121 | 0.5973 | 0.571 | 0.571 |

**Table 5**. One-sample Wilcoxon tests comparing the change in tapping asynchrony in each condition and time point to zero for Experiments 5 and 6.

|  | Distractor Type | Time Point | N | W | z | p | p (corrected) |
| --- | --- | --- | --- | --- | --- | --- | --- |
| Experiment 5 (online) | Large pitch shift | 250 | 52 | 640 | -0.446 | 0.659 | 0.902 |
|  |  | 750 | 52 | 352 | -3.069 | 0.002 | 0.017 |
|  |  | 1250 | 52 | 675 | -0.128 | 0.902 | 0.902 |
|  |  | 1750 | 52 | 618 | -0.647 | 0.521 | 0.902 |
|  | Small pitch shift | 250 | 52 | 585 | -0.947 | 0.346 | 0.902 |
|  |  | 750 | 52 | 717 | 0.255 | 0.802 | 0.902 |
|  |  | 1250 | 52 | 652 | -0.337 | 0.740 | 0.902 |
|  |  | 1750 | 52 | 611 | -0.710 | 0.480 | 0.902 |
| Experiment 6 (in-lab) | Large pitch shift | 250 | 21 | 122 | 0.226 | 0.838 | 0.973 |
|  |  | 750 | 21 | 44 | -2.485 | 0.011 | 0.045 |
|  |  | 1250 | 21 | 202 | 3.007 | 0.002 | 0.013 |
|  |  | 1750 | 21 | 125 | 0.330 | 0.759 | 0.973 |
|  | Small pitch shift | 250 | 21 | 78 | -1.303 | 0.203 | 0.541 |
|  |  | 750 | 21 | 117 | 0.052 | 0.973 | 0.973 |
|  |  | 1250 | 21 | 119 | 0.122 | 0.917 | 0.973 |
|  |  | 1750 | 21 | 100 | -0.539 | 0.609 | 0.973 |

**Table 6**. Wilcoxon signed rank tests comparing the change in tapping asynchrony between large and small pitch shift at each time point for Experiments 5 and 6.

|  | Time Point | N | W | z | p | p (corrected) |
| --- | --- | --- | --- | --- | --- | --- |
| Experiment 5 (online) | 250 | 52 | 730 | 0.373 | 0.712 | 0.873 |
|  | 750 | 52 | 328 | -3.288 | 0.001 | 0.004 |
|  | 1250 | 52 | 779 | 0.820 | 0.415 | 0.830 |
|  | 1750 | 52 | 671 | -0.164 | 0.873 | 0.873 |
| Experiment 6 (in-lab) | 250 | 21 | 134 | 0.643 | 0.539 | 0.719 |
|  | 750 | 21 | 32 | -2.902 | 0.002 | 0.010 |
|  | 1250 | 21 | 181 | 2.277 | 0.022 | 0.043 |
|  | 1750 | 21 | 126 | 0.365 | 0.733 | 0.733 |
